# Correlation of invitro susceptibility based on MICs and SQLE mutations with clinical response to terbinafine in patients with tinea corporis/cruris

**DOI:** 10.1101/326603

**Authors:** Ananta Khurana, Aradhana Masih, Anuradha Chowdhary, Kabir Sardana, Sagar Borker, Aastha Gupta, R K Gautam, P K Sharma, Dhruv Jain

**Author notes:** Address correspondence to Ananta Khurana, and Anuradha Chowdhary.

## Abstract

Recalcitrant dermatophytoses are on the rise and recent publications have documented high minimum inhibitory concentrations (MICs) to TRB and squalene epoxidase (SQLE) mutations. However, literature correlating the laboratory the data with clinical response is lacking.

This study was conducted to study the clinico-mycological profile of tinea corporis and cruris, including antifungal susceptibility testing (AFST) and SQLE mutation analysis and correlate these with clinical response to TRB. Skin scrapings of patients with tinea corporis with/without tinea cruris were subjected to species identification, AFST and SQLE gene analysis (on 15 isolates). KOH confirmed cases were started on TRB 250mg once a day (OD). If >50% clinical clearance was achieved by 3 weeks; the same dose was continued.(Group 1). If clinical clearance at 3 weeks was <50%, the dose was increased to 250mg twice a day (BD) (Group 2). If the response still remained below 50% after 3 weeks of BD, the patients were treated with itraconazole (ITR)(Group 3). *Trichophyton interdigitale* was confirmed on all 64 isolates obtained on culture. Forty four (68.7%) isolates had high (≥1 μg/ml) MICs to TRB. Six isolates were found to have aminoacid substitution Leu393Phe in SQLE protein, while one had the substitution Phe397Leu. The difference in modal MICs to TRB between the 3 clinical response groups (1.5157μg/ml, 5.0396 μg/ml and 20.1587μg/ml respectively for group 1,2 and 3) was highly significant. Clinical response was achieved in 68% of those resistant by MIC data, and 42.8% of SQLE mutation harboring isolates, by increasing drug (TRB) exposure.

We infer that TRB resistance in dermatophytes has reached alarming proportions in our patients. Though improved outcomes were achieved with higher drug exposure, with the high failure rate seen in the study, the case for shifting to another class of antifungals as first line agent against dermatophytoses is strong.

## INTRODUCTION

Dermatophytoses of skin have been successfully managed with terbinafine (TRB) in the past. The standard recommendation for tinea corporis/cruris has been TRB 250mg OD for 2-3 weeks; and in fact even lesser treatment durations have been reported to be successful. (1–3) Of late though, there has been an upsurge in difficult to treat dermatophytoses and declining responses to TRB have been on record.(4–7) High minimum inhibitory concentrations (MICs) and mutations in the target enzyme SQLE have been demonstrated in a few recent reports and the use of higher TRB dosages is becoming common.(6,8–13) However, clinical breakpoints have not been defined for TRB in dermatophytoses and the in vitro data cannot be directly applied to clinical situations.

With this work, we aimed to study the clinico-mycological profile of our patients with corporis/cruris and correlate the clinical response achieved with the standard treatment and higher dose/durations of TRB with the MICs obtained in laboratory and with target gene (SQLE) mutations. We also analysed the response of TRB failures to ITR.

## PATIENTS AND METHODS (11)

The study was approved by the institutional ethics committee and registered with the clinical trials registry of India (CTRI). The presented patients were recruited between July 2016 to December 2017. Eighty five consecutive patients of tinea corporis, with or without tinea cruris, presenting to the dermatology out patients department of Dr Ram Manohar Lohia hospital were included after obtaining an informed consent. Those with co-existent tinea manuum/pedis/capitis or tinea unguium were excluded. Diagnosis was made on clinical examination and confirmed by KOH microscopy. The included patients had at least 5 lesions over the glabrous skin and/or large lesion/s covering significant body surface area, and judged by the primary investigator (AK) as requiring systemic treatment. Patients who had used any systemic anti-fungal in the preceding 4 weeks or used any topical antifungal or steroid in the preceding 2 weeks were excluded. Pregnant or lactating women and children less than 12 years of age and/or weighing less than 40 kg were also excluded. A detailed history of disease onset, duration, course, family history and previous treatments was taken followed by examination of the entire skin surface to look for lesions. Photographs (with Canon Powershot G12) were taken for documentation and comparison of treatment response. The scales collected were transported to the Medical Mycology laboratory of Vallabhbhai Patel Chest Institute, Delhi, in a thick dry sheet of paper. Treatment was started after sending the samples and with KOH confirmation. Complete cure was defined as complete clinical clearance along with a negative KOH from the site of initial sampling.

The patients were started on terbinafine in a dosage of 250 mg OD and asked to follow up 3 weekly. The treatment response was judged by the primary investigator at each visit by clinical examination (based on extent of lesions, erythema and scaling) and comparison with previous photographs. If the response to this regimen was greater than 50% at the 3 week follow up, the same dose was continued till complete cure (Group 1). If the clinical response after 3 weeks was less than 50% or if new lesions appeared during this time, the patient was shifted to terbinafine 250mg BD, and reassessed after another 3 weeks. If >50% clinical clearance was achieved by this point, the same regimen was continued till complete cure (Group 2). However, if the response still remained below 50%/new lesions appeared, the patient was shifted to ITR given in a dose of 100mg BD and treated till complete cure. (Group 3) We chose a cutoff of 3 weeks as this has been the standard recommended treatment duration for tinea corporis/cruris.(1,14) The patients were counseled on general hygiene measures and ways to reduce transmission among family members. No topical antifungal was given to avoid additive effect and patients were instructed to use only emollients topically. Antihistamines were prescribed as per the patient’s symptoms. Patients were asked of adverse effects pertaining to TRB at each visit and liver function tests were performed periodically.

Skin scrapings were processed for direct microscopic examination by 10% potassium hydroxide (KOH)/Blankophor. The specimens were inoculated on two sets each of Sabourauds dextrose agar (SDA) containing gentamicin and chloramphenicol and the other containing cycloheximide (0.05%) incubated at 28°C for 2 weeks. Preliminary macroscopic phenotypic identification was done on potato dextrose agar (PDA) incubated at 28°C for 14 days. Slide cultures prepared on PDA and mounted in lactophenol cotton blue were examined microscopically.

Molecular identification of isolates was performed by sequencing the internal transcribed spacer (ITS) region of the small subunit ribosomal deoxyribonucleic acid (rDNA). ITS sequences were subjected to BLAST searches at GenBank (http://www.ncbi.nlm.nih.gov/BLAST/Blast.cgi). Sequence-based species identification was defined by ≥99% sequence similarity with ≥99% query coverage. The neotype and type strains of *Trichophyton* species (*T. interdigitale*, CBS 428.63NT; *T. mentagrophytes*, CBS 318.56NT146;*T. rubrum*, CBS 392.58NT; *T. tonsurans*, CBS 496.48NT and *T. violaceum*, CBS374.92T147) were retrieved from GenBank and included for phylogenetic analysis.

Antifungal susceptibility testing (AFST) was carried out using the Clinical and Laboratory Standards Institute broth microdilution method (CLSI-BMD), using the M38-A2 guidelines. Eleven systemically/topically used antifungals were tested including terbinafine (TRB, Synergene India, Hyderabad, India,), itraconazole (ITR; Lee Pharma,Hyderabad, India), voriconazole (VRC; Pfizer Central Research, Sandwich, Kent, UnitedKingdom), fluconazole (FLU; Sigma-Aldrich, Germany),sertaconazole (SER; Optimus, Hyderabad, India), luliconazole (LUZ, Sun pharmaceuticals, Baddi, HP, India), clotrimazole (CLT; Sigma-Aldrich), miconazole (MCZ; Sigma-Aldrich), ketoconazole (KTC; Sigma-Aldrich), amphotericin B (AMB; Sigma-Aldrich) and griseofulvin (GRE; Sigma-Aldrich). CLSI recommended control strains of *Candida krusei*ATCC6258 and *Candida parapsilosis*ATCC22019 were included for every batch of isolates tested each day. Reference strains of *T. interdigitale*(ATCC MYA-4439) and *T. rubrum* (CBS 592.68) were also included in susceptibility testing. Minimum inhibitory concentration endpoints for all the drugs were defined as the lowest concentration that produced complete inhibition of growth as read visually at 72h.

The amplification primers for *SQLE* gene, as described previously were used.^10^ PCR was carried out in a 50 μL reaction volume and the conditions included initial denaturation for 5 minutes at 95°C followed by 34 cycles of 30 seconds at 95°C, 30 seconds at 60°C and 180 seconds at72°C. DNA sequencing was performed using the PCR primers at2.5 mmol/L concentration. All sequencing reactions were carried out in a 10 μL reaction volume using BigDye Terminator Kit v3.1(Applied Biosystems, Foster City, CA, USA) according to the manufacturer’srecommendations and analysed on an ABI3130xL GeneticAnalyzer (Applied Biosystems). The amino acid sequence of SQLEp of all the investigated *T. interdigitale* in the present study was compared with reference sequence of *T. interdigitale* (GenBank accession number EZF33561).

Statistical analysis was done on SPSS software version 21. One way ANOVA was used to compare the geometric mean (GM) MICs between the 3 treatment response groups. Chi square test was used to compare the treatment response with demographic data.

## RESULTS

A total of 85 patients were included. Of these, 64 samples (75.3%) could be grown on culture and were taken up for further analysis. Out of these 64 culture positive patients, there were 44 males and 20 females. Ages of patients ranged from 14 to 71 years, with a mean age of 34.9 years. Disease duration ranged between 3 weeks to 5 years, with an average duration of 8.8 months. Thirty five (54.6%) patients had a positive family history of dermatophytic infections, either at present or in recent past. Interestingly, 45 patients gave a history of sharing of towels between family members, bringing out the importance of fomite transmission in households. Only 2 had pets at home and those were apparently uninfected as per the patients’ account. Eight were involved in regular outdoor sports activities and 37 gave history of prolonged working in hot and humid conditions, either at home or as a part of their vocation.

Most patients (51) gave a visual analogue score (VAS) between 8 to 10 for associated pruritus. Forty five (70.3%) patients had a history of application of topical steroid creams, either plain steroid or as a combination with antifungals. The most commonly used topical steroid was clobetasol (27) followed by betamethasone (17), beclomethasone (5), fluticasone (2) and mometasone (2). Seven had used more than one class of topical steroid. Most (32; 71.1%) had procured these over the counter from pharmacists without a prescription, while the rest (13, 28.8%) were prescribed by general practitioners. Nine had apparent signs of topical steroid abuse in the form of striae, skin thinning and depigmentation. Three of the patients were diabetics on treatment, while one each had coronary artery disease and iron deficiency anemia.

Most patients (30; 46.8%) had 5-10 lesions at presentation, while 16 (25%) had 10-20, 13 (20.3%) had more than 20 and 5(7.8%) had 5 or less lesions. Thirty eight (59.3%) had large confluent lesions covering extensive areas. Associated inflammation on clinical examination was moderate in most (34;53.1%) cases, mild in 20 (31.25%) and severe with pustulation in 3 (4.6%). Seven (10.9%) had mixed picture with varying degrees of inflammation at different body sites. *Trichophyton interdigitale* was isolated from all 64 patients. The MICs for terbinafine, itraconazole, fluconazole, voriconazole, ketoconazole, amphotericin B, griseofulvin, miconazole, clotrimazole, luliconazole and sertaconazole are tabulated in Table 1. Luliconazole had lowest MICs (MIC50 0.0007μg/ml) while MICs for fluconazole were consistently high (MIC50 32μg/ml). Forty four(68.75%) isolates had high MICs against TRB (MIC ≥1 μg/ml) and 23 isolates (35.9%) had extremely high MICs of ≥32μg/ml. Nineteen (29.6%) isolates had MIC of 0.5μg/ml while only 1 had MIC of 0.25μg/ml, the lowest recorded in our patients.

**TABLE 1:** MICs of the 56 isolates to the 11 antifungals tested.

| **Parameter**<br>($\mu\text{g/ml}$) | **TRB**<br>($\mu\text{g/ml}$) | **ITR**<br>($\mu\text{g/ml}$) | **FLU**<br>($\mu\text{g/ml}$) | **VRC**<br>($\mu\text{g/ml}$) | **KTC**<br>($\mu\text{g/ml}$) | **AMB**<br>($\mu\text{g/ml}$) | **GRI**<br>($\mu\text{g/ml}$) | **MCZ**<br>($\mu\text{g/ml}$) | **CLT**<br>($\mu\text{g/ml}$) | **LUZ**<br>($\mu\text{g/ml}$) | **SER**<br>($\mu\text{g/ml}$) |
| --- | --- | --- | --- | --- | --- | --- | --- | --- | --- | --- | --- |
| **GM** | 2.89 | 0.414 | 16.53 | 0.285 | 0.878 | 0.374 | 3.75 | 2.327 | 2.327 | 0.007 | 1.176 |
| **MIC<sub>50</sub>** | 1 | 0.5 | 32 | 0.25 | 1 | 0.38 | 4 | 2 | 2 | 0.007 | 1 |
| **MIC<sub>90</sub>** | 32 | 1 | 64 | 1 | 2 | 1 | 8 | 4 | 4 | 0.015 | 6.4 |
| **RANGE** | 0.25-<br>$\geq 32$ | 0.06-<br>$\geq 16$ | 0.5-<br>$\geq 64$ | 0.06-2 | 0.25-<br>$\geq 32$ | 0.25-1 | 0.5-<br>$\geq 8$ | 0.25-<br>$\geq 16$ | 0.5-8 | 0.0035-0.125 | 0.25-<br>$\geq 16$ |

GM: Geometric mean, TRB: Terbinafine, ITR: Itraconazole, FLU: Fluconazole, VRC:
Voriconazole, KTC: Ketoconazole, AMB: Amphotericin B, GRIS: Griseofulvin, MCZ:
Miconazole, CLT: Clotrimazole, LUZ: Luliconazole, SER: Sertaconazole

Among the culture positive patients, complete follow up data was available for 30 patients and these were further included for the clinico-mycological correlation. (Table 2) Fifteen (50%) out of these responded to prolonged duration of OD TRB, given beyond 3 weeks in all but 2 patients. The mean duration of response in this group was 39.46±13.12 days (range 21-66 days). Out of these 15, 8 had MIC of 1μg/ml, 2 had MIC of 0.5 μg/ml, 1 each had MIC of 0.25μg/ml,2μg/ml and 4μg/ml and 2 had MIC of ≥32μg/ml. The GM MICs to TRB in this group was 1.515μg/ml. Only one patient in this group had history of previous terbinafine exposure, while 10 (66.67%) had applied topical steroids over their lesions previously.

**TABLE 2:**
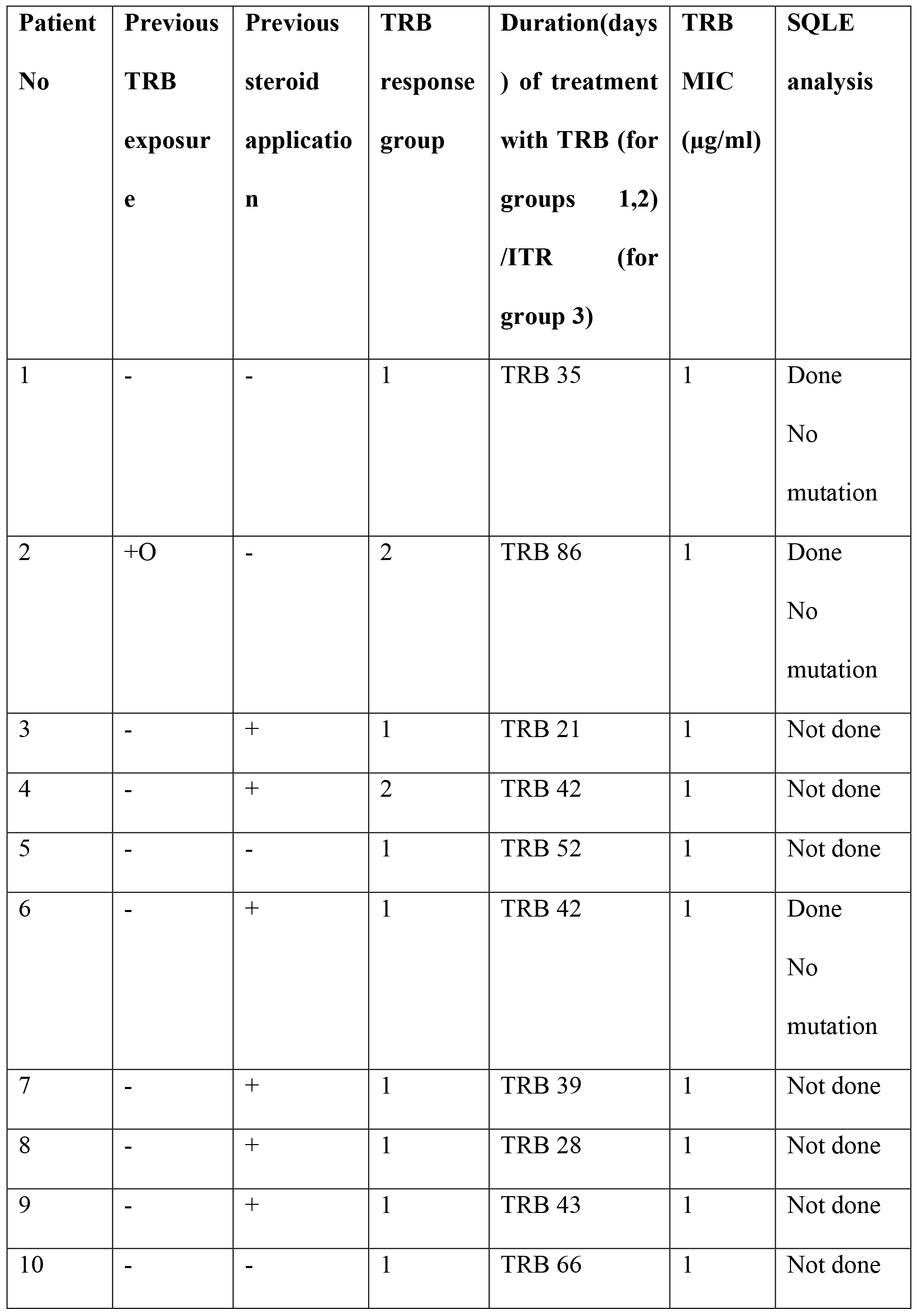
Comparison of clinical response and mycological data of 30 patients.

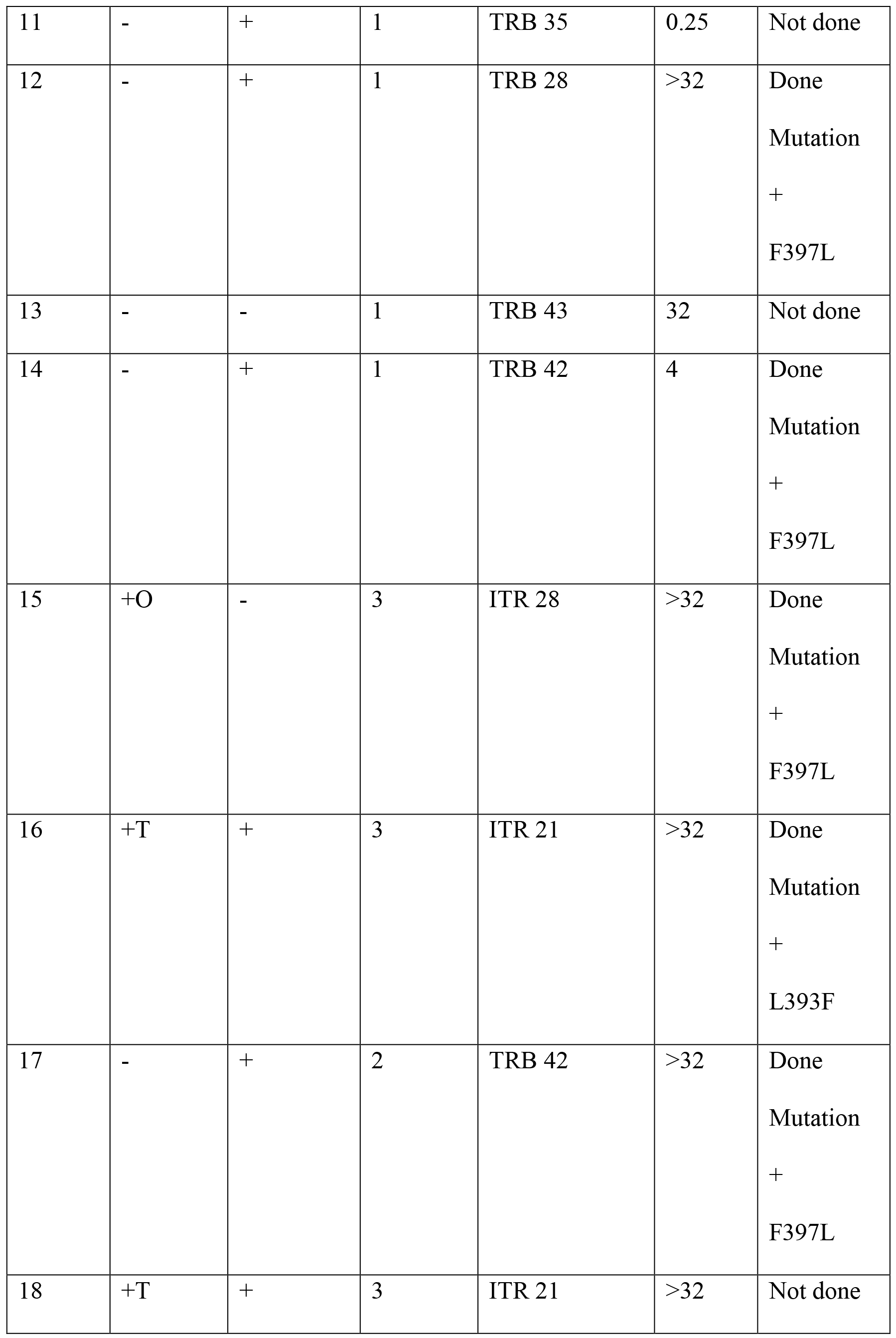

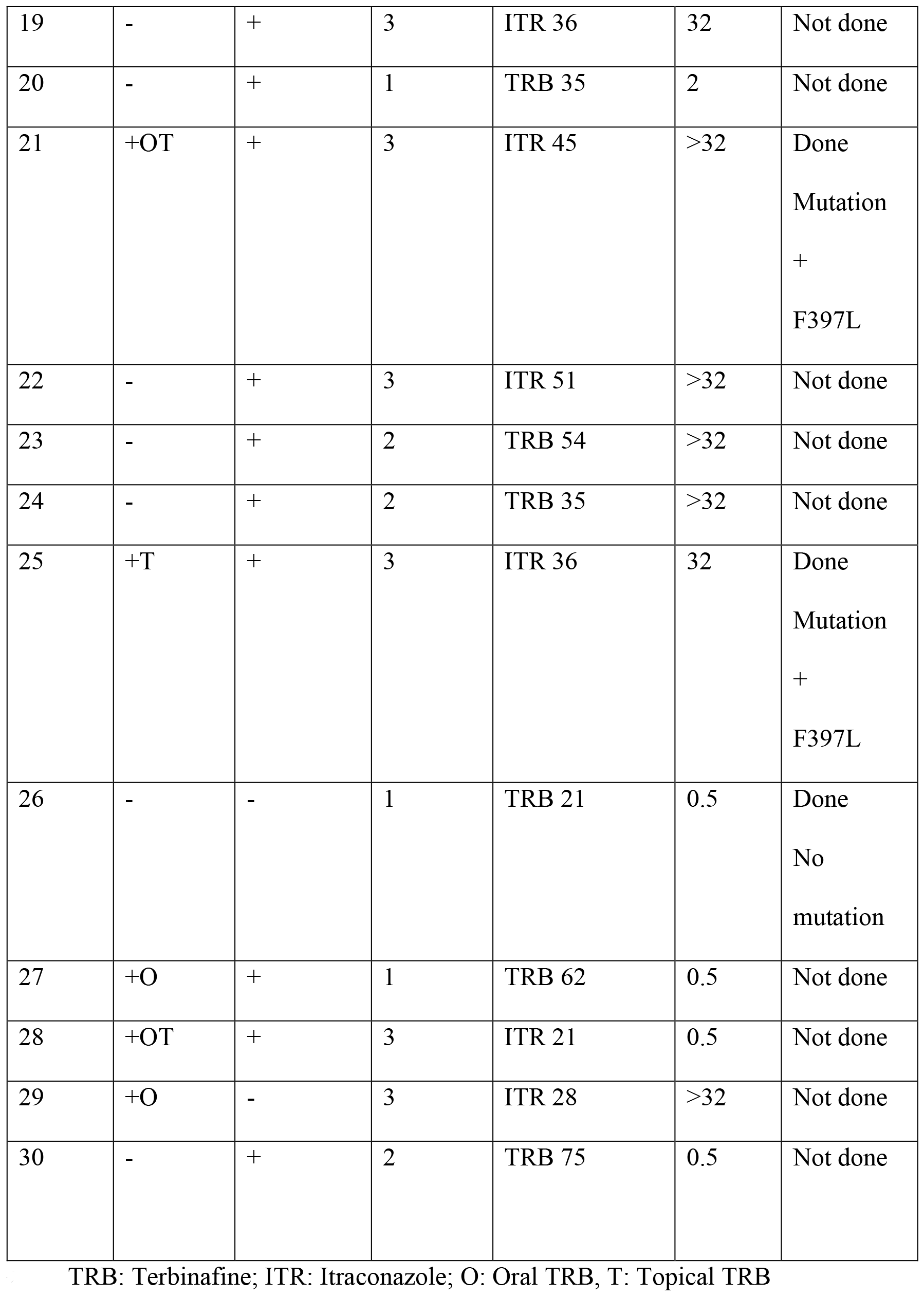

The other 15 patients did not show a clinically relevant response (>50% clearance) to OD TRB till 3 weeks and were shifted to BD dosing. Six of these showed >50% clearance after 3 weeks of updosing and were continued on this dose till complete cure. The TRB MIC distribution in this group was 1μg/ml in 2, ≥32μg/ml in 3 patients and 0.5μg/ml in 1. The GM MIC for the group was 5.039μg/ml and average duration of treatment (including OD and BD treatment) was 55.66±20.48 days (range 42-86 days). Only one patient in this group had used terbinafine previously, while 5 had applied topical steroids previously.

Finally, 9 patients did not achieve 50% clinical clearance even after 6 weeks of TRB (3 weeks of OD and 3 weeks of BD) and were treated with itraconazole(ITR) 100 mg BD for duration ranging from 21 to 51 days (mean 31.88±3.64 days). Eight of these had TRB MICs of ≥32μg/ml, while one had MIC of 0.5μg/ml. The GM MIC to TRB was 20.158μg/ml. Seven of these had been treated with TRB previously (2 with oral TRB, 3 with topical TRB and 2 with both oral and topical TRB) and 7 had applied topical steroids previously. MICs to ITR in the 9 patients ranged from 0.125 to 1μg/ml, with GM MIC of 0.314μg/ml.

The difference between the GM MICs between the three groups was highly significant (p=0.004), as was the difference between GM MICs of combined group 1 and 2 (patients who achieved cured with TRB, with any dose/duration; 2.136μg/ml) and group 3 (who were not cured even with higher drug exposure; 20.159μg/ml) (p=0.004). However, we didnot find a statistically significant correlation between individual MICs or susceptibility based on MIC (MIC≥1 μg/ml or <1 μg/ml) with cure achieved with TRB (with standard or higher drug exposure). We also did not find any significant association between cure achieved with TRB or MIC values to TRB with family history of tinea, duration of the disease, past history of topical steroid use or history of episodes of tinea in recent past.

Among those infected with TRB susceptible organisms (MIC<1 μg/ml), 80% achieved cure, while 68% of those infected with resistant organisms achieved cure, either by standard dose/duration of treatment or by updosing/increasing treatment duration. (Table 3)The odds of achieving cure with TRB, when infected with a susceptible organism were 1.88 times the odds of achieving cure when the organism was TRB resistant. Further, of the 4 susceptible TRB responsive patients, one achieved cure with standard dose and duration of TRB, 2 to longer duration of the standard dose and 1 after updosing to BD. Of those infected with resistant organism, most of those who responded (11), responded with an increase in the treatment duration of OD dose, while 5 achieved cure with updosing.

**TABLE 3:** Response to TRB in mycologically susceptible and resistant infections.

|  | **Organism susceptible**<br>**(MIC<1µg/ml); n=5** | **Organism resistant**<br>**(MIC>1 µg/ml),**<br>**n=25** | **Odd's ratio** |
| --- | --- | --- | --- |
| **Cure achieved with**<br>**TRB; n=21** | 4(80%) | 17 (68%) | 1.88 |
| **Not cured with TRB**<br>**N=9** | 1(20%) | 8 (32%) |  |

Squalene epoxidase (SQLE) gene mutation analysis was done in a total of 15 (out of 64) isolates. Out of these, 7 harbored mutations leading to single amino acid substitution in the SQLE protein. (TABLE 2) The MICs in mutated isolates were 4 μg/ml (1 isolate) and >32 μg/ml (6 isolates)The 8 isolates which did not demonstrate mutations had MICs of 0.5 μg/ml (5) and 1 μg/ml (3). Six of the mutation harboring isolates had the aminoacid substitution Leu393Phe in SQLE protein and one had the substitution Phe397Leu. Four of these belonged to treatment group 3, 1 to group 2 and 2 to group 1. Four (57.1%) of these patients had previous exposure to TRB (oral/topical). Three patients (42.8%) with SQLE mutations responded to higher dose/longer duration of treatment, while 4 (57.14%) did not respond to the drug even after increasing the drug exposure.

## DISCUSSION

A predominant finding of our study is the very high level (68.7%) of TER resistance in our isolates. Secondly, correlation of the clinical response to TRB with mycological parameters of MIC and SQLE gene mutations revealed that increasing TRB drug exposure is able to achieve cure in about 68% of isolates resistant by MIC data and about 57% of confirmed SQLE mutated isolates’ infections. The data we have presented here forms the basis of establishing clinical breakpoints for a drug-species pair, although larger patient numbers and intricate tissue level PK data would also be essential for the same.

Dermatophytoses present a huge economic burden on the medical establishments all over the world, with an estimated worldwide prevalence of 20-25%.(15) However, the drug classes available against dermatophytes are limited in their spectrum of action, largely targeting the ergosterol biosynthetic pathway. Terbinafine has been the drug of choice and has been in use for almost three decades now. However, there are still lacunae in our knowledge regarding some aspects of its use for dermatophytoses mainly relating to the pharmacokinetic/pharmadynamic (PK/PD) parameter best predictive of response and the clinical breakpoints.

It has been previously demonstrated in in-vitro trials that the frequency of naturally occurring mutants with resistance to TRB and of development of resistance during prolonged exposure to the drug are both very low (~ 10^−9^), compatible with the reported mechanism of single nonsilent nucleotide substitution in gene encoding SQLE protein.(16–17)Indeed, prior to 2017, there had been only 2 documented cases of TRB resistance in dermatophytes.(18–19) However, the scenario has been increasingly reported over the last year. (Table 4) Apart from SQLE mutations, there are isolated reports of TRB resistance mediated by mutations in salA gene, encoding salicylate1-monooxygenase and genes encoding ABC transporter proteins.(21–22)

**TABLE 4:**
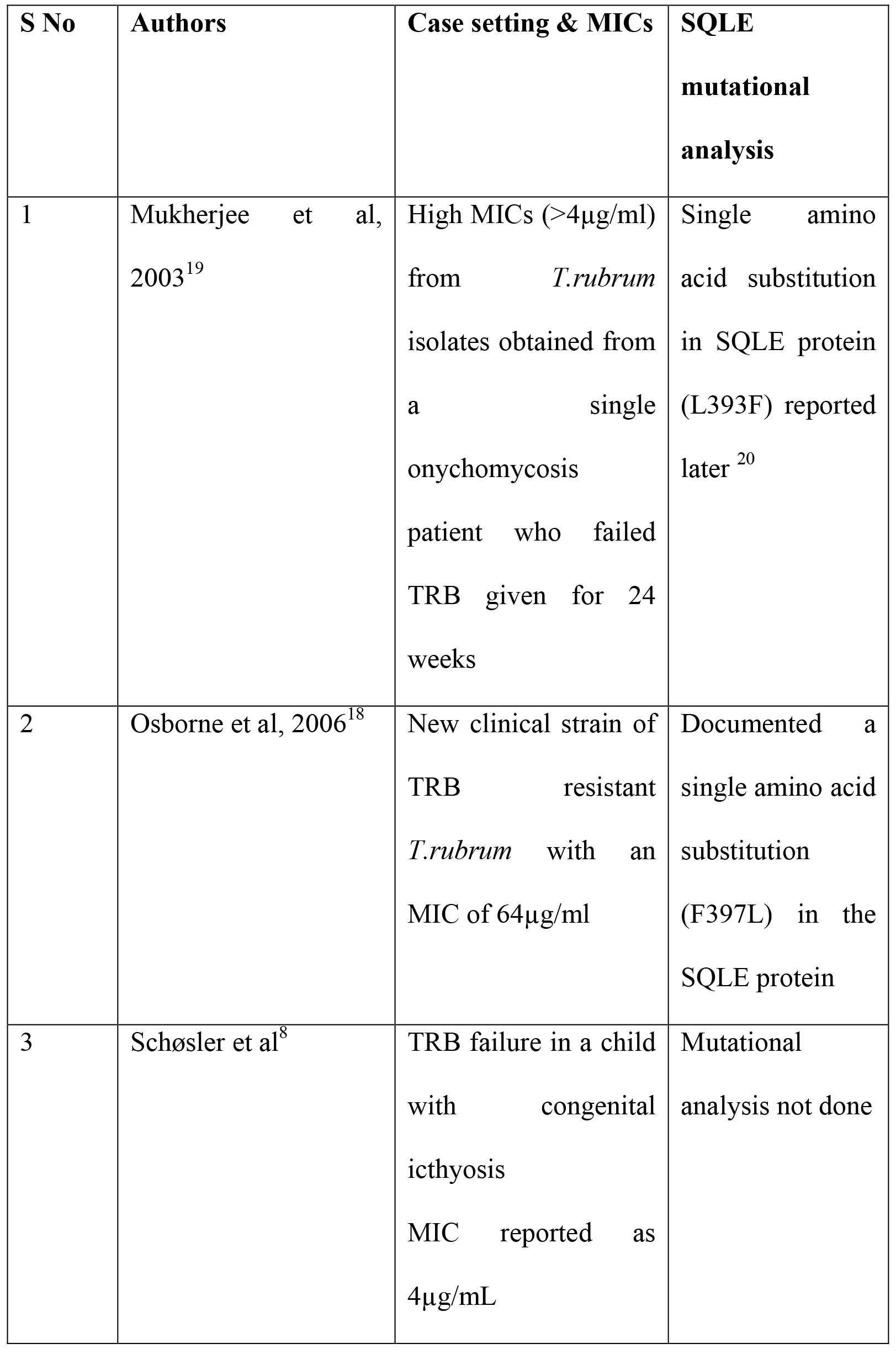
Previous reports of TRB resistance in dermatophytes.

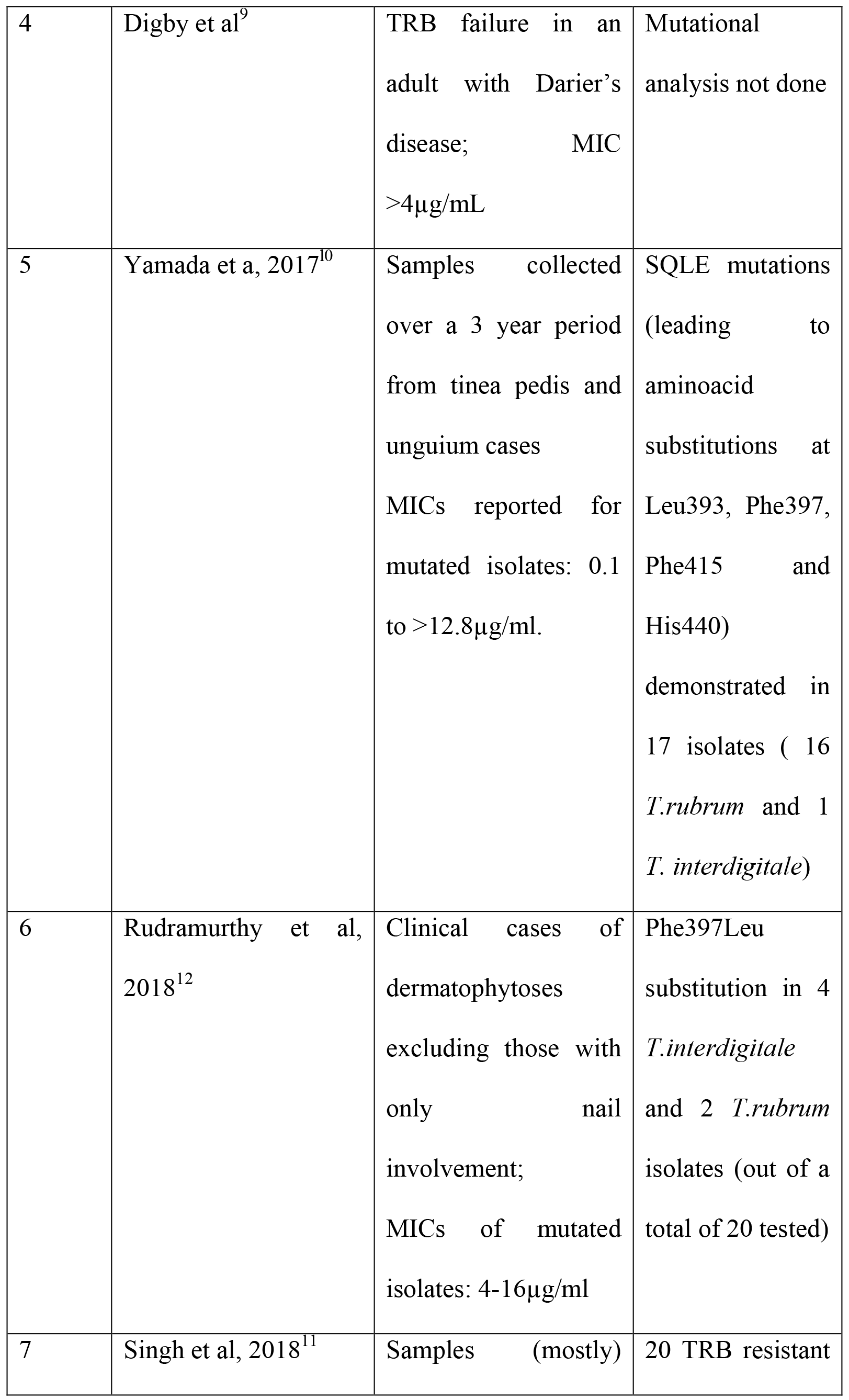

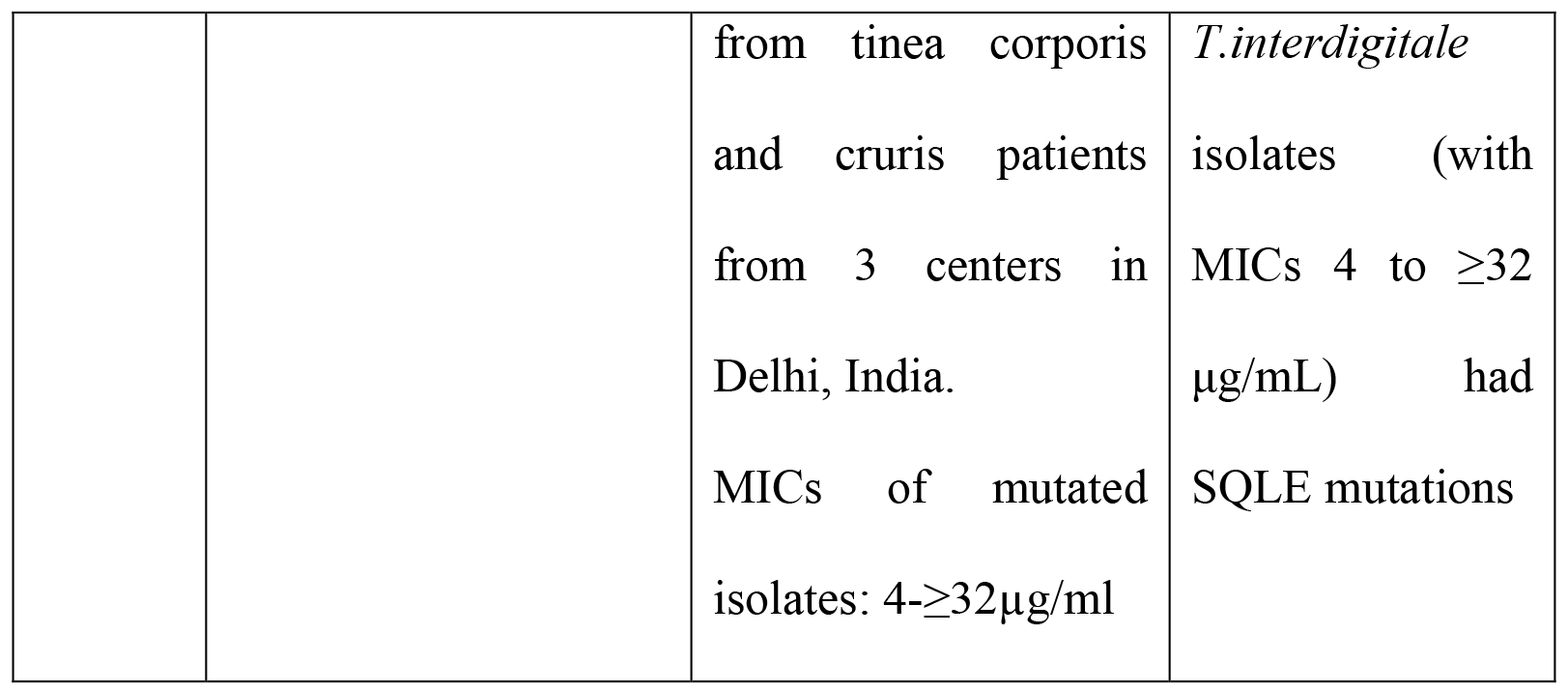

An important lacuna in literature related to the topic at this point of time is the lack of prospectively acquired clinical correlation data. As in vitro data may not be directly extrapolatable to clinical situations, the clinical correlation in individual cases becomes an important domain to explore. Further, it is also important to assess host factors/trends which may possibly be contributing to development of resistance to TRB, considered an unusual event so far.

A striking finding of our study is the complete dominance of *T.interdigitale* as the etiological agent for tinea corporis/cruris. Literature from most other nations as well as older Indian literature mostly cites *T.rubrum* as the predominant organism, and *T.interdigitale* has previously been reported as a prominent species from only a few geographical areas.(23–28) However, few other recent Indian reports have also found *T.interdigitale* in a large percentage of isolates.(11,12,29) A species shift has been considered as an important factor for the epidemic of recalcitrant tinea India is facing since few years, although the reasons for it are not yet clear.

Our results also reconfirm the unfortunate trend of high MICs to TRB. In all, 68.75% of the 64 isolates had MICs of ≥1μg/ml, with 35.9% having very high MICs of >32μg/ml. No epidemiologic cut offs (ECVs) have been established for terbinafine resistance in *T. interdigitale* by CLSI. We have taken the cutoff as described for *T.rubrum* strains previously.(30)The criteria of high TRB MICs of ≥1 μg/ml however needs to be validated for *T.interdigitale* by multi-centric studies with large number of isolates using the defined reference broth microdilution method. Fluconazole resistance (MIC≥64μg/ml), was seen in 5 (8.9%) isolates.(31)Griseofulvin has become largely ineffectual for dermatophytoses over years and the same trend was reported in our study as well, with most isolates showing high MICs (GM MIC of 3.75μg/ml).(32)The lowest MICs were seen with luliconazole (GM MIC of 0.007μg/ml). Voriconazole (GM MIC 0.285μg/ml), itraconazole (GM MIC 0.414μg/ml), ketoconazole (GM MIC 0.878μg/ml) and amphotericin B (GM MIC 0.374μg/ml) also demonstrated low MICs. Only 1 isolate was resistant to itraconazole (MIC≥8μg/ml), 1 to voriconazole (MIC≥2μg/ml) and 3 to ketoconazole(MIC≥8μg/ml).(31) The topicals miconazole and clotrimazole, widely used for dermatophytoses as OTC as well as prescription drugs, also showed low activity with GM MICs of 2.327μg/ml. Overall the MICs reported in our study were higher than those previously reported with dermatophytes for most drugs, though the difference in TRB values was most striking.(32–35)

It was interesting to note that only two (Patient 3, TRB MIC 1μg/ml and patient 26, TRB MIC 0.5μg/ml) of the 30 analyzed patients responded to the conventional dose and duration of TRB. The rest in group 1 required much longer courses ranging from 28 to 66 days. Notably, all but 3 of the group 1 patients had MICs of 1μg/ml or more, but responded to longer courses of the drug. Extrapolating from the data on azoles, it is known that optimizing drug exposures can improve clinical outcomes in the wake of raised MICs/resistant strains, depending on the variables like the degree of rise in MICs, resistance mechanism and pharmacokinetic/pharmacodynamics (PK/PD) properties of the antifungal agent.(36)For drugs used against dermatophytoses, the levels in tissue (stratum corneum [SC]) are more important than the plasma levels. But, there is only scant literature on tissue level PK of TRB. From 2 such studies done by Faergemann et al, it can be interpreted that longer courses lead to higher levels of TRB in the SC. The authors reported SC levels of 14.4μg/g after administration of TRB 250mg OD for 28 days vs 7.63μg/g at the end of a 14 day course and 2.52μg/g at the end of a 7 day course of the same dose.(37–38) The higher levels thus achieved with longer durations may have been able to surmount the higher MICs in group 1 patients.

Further, on updosing to BD, another 6 patients achieved complete cure. As defined in previously done PK studies, TRB has a linear PK profile upto750mg of dose, implying that an increase in dose till this level would increase plasma levels proportionately.(39) And this would in all likelihood lead to an increased SC level, unless the uptake by skin is saturable, of which there is no documentation yet to the best of our knowledge. This group benefitted both from an increase in dose and an increase in duration, possibly achieving higher SC levels than achieved by group 1. Notably, the GM MIC of group 2(5.039μg/ml) was about 5 times higher than that of group1 (1.5157μg/ml). This may explain why the group 2 patients did not respond only with longer durations of treatment.

Finally 9 patients did not achieve 50% clinical clearance even after 6 weeks of TRB (OD for 3 weeks followed by BD for 3 weeks) and were then treated with itraconazole. The GM MIC of this group was 20.159μg/ml, which was about 20 times that of group 1 and 4 times that of group 2. This demonstrates a trend wherein higher drug exposure surmounts the higher MICs to an extent, after which the drug fails to work. Whether doses higher than what we used would improve cure rates further may be an area of further research, though further increases may be limited by drug toxicity. We did not observe any drug related adverse effects with using longer durations and higher doses of TRB, although the data may be biased by a small sample size.(40) Analyzing the clinical response data of the 7 patients in whom SQLE mutations were characterized, 4 of these (57.1%) did not respond to higher drug exposure, while 3 (42.8%) did. This data, though small, largely supports the 90-60 rule of clinical correlation with invitro susceptibility testing, wherein about 60% of resistant isolates respond to the drug in vivo.(41) The susceptibility based on MIC data also gives a similar interpretation, wherein 68% of the isolates with MIC≥1 μg/ml could be successfully treated with TRB. One of these (patient 3) responded to the usual dose and duration of TRB, while the other 16 required longer durations/higher dose. In both the present study and our previous work, SQLE mutations were noted in isolates with MICs 4 to ≥32 μg/mL and wild type genotype (i.e., no SQLE mutation) was noted in *T. interdigitale* isolates with MICs ≤2 μg/mL This may imply that SQLE mutations lead to a high level resistance in*T. interdigitale* and alternate mechanisms may be working in resistant isolates with MICs <4 μg/mL.(11)

We couldnot identify any host factors predisposing to TRB resistance. Topical steroid use may enhance development of resistance to antifungals being used simultaneously, by activating fungal metabolism and by a cell membrane protective activity. However, we did not find a statistically significant correlation between topical steroid use and clinical response to TRB. One possible reason for this could have been the widely prevalent use of topical steroids in our patients which precluded formation of comparable groups. Indeed, 21 of the 30 analyzed patients had used topical steroids before, while only 9 had not. Prior TRB exposure also did not yield any predictable results although on general observation of the trend among the 3 treatment groups, we observed previous TRB exposure to be most prominent in group 3 (77.78% in group 3 vs 6.67% in group 1 and 16.67% in group 2). Age, family history of tinea and history of recurrent tinea in the past also had no significant correlation with TRB resistance (based on MIC values). Whether the resistance we have seen was primary or acquired cannot be commented upon a single time assessment. But it was an interesting finding that most (16/17; 94.11%) of those who were resistant and yet responded to TRB had not been exposed to TRB before, while most of those who were resistant and did not respond to TRB (7/8; 87.5%) had been exposed to TRB before.

All 9 patients who failed TRB were successfully treated with ITR. These all were sensitive to ITR based on MIC data as well. Overall too, ITR resistance was seen in only one of the 64 isolates. Thus, ITR sensitivity seems to be preserved in dermatophytes so far, and ITR may become a frontline drug for dermatophytoses with rising failures being seen with TRB.

In the end, we would like to highlight that resistance to TRB has reached a worrisome level in isolates from dermatophyte infections in our patients. Increasing drug exposure by means of higher dose/duration can surmount this to some degree, but the high failure rate still (30%) cannot be ignored. Although localized to a few geographical locales so far, it may not be long before the resistance spreads to other regions and it would be desirable to preplan strategies to combat the same. A rethink on the treatment order and development of newer drug classes against dermatophytes with novel mechanisms of action is due.

